# The effect of basal core promoter and pre-core mutations on HBV replication and persistence in mice

**DOI:** 10.1101/2020.10.26.354944

**Authors:** Yi Chen, Zhiwei Xu, Yanli Zeng, Junping Liu, Yongjun Tian, Yi Kang

## Abstract

The appearance of the BCP or Pre-C mutations, which reduce or abolish HBeAg production, could increase HBV replication. The remove of the HBeAg often lead to a vigorous immune response, which has an important role in HBV related fulminant outcome. In this study, BCP mutations and Pre-C mutations were separately introduced by site-directed mutagenesis in the same genetic background of an HBV infectious clone, to determine the effect of these mutations per se on replication. BCP and Pre-C mutations increased HBV replication both in *vitro* and *in vivo*. HBV could persist in mice injected with wild type HBV infectious clone for about 7 weeks. However, HBV could persist about 5 weeks in mice injected with BCP HBV infectious clone, and 3 weeks only in mice injected with Pre-C HBV infectious clone. HBV related CD8+ CTL response in BCP HBV infectious clone injected mice only slightly increased, but significantly increased in Pre-C HBV infectious clone injected mice compared with that in wild type HBV infectious clone injected mice. The population of Tregs significantly increased in liver but not in spleen of mice injected with Pre-C HBV infectious clone. In summary, we demonstrate that HBeAg plays an important role in suppressing the CTL response, which is related with increasing the frequency of Tregs in mouse. Lack of HBeAg expression leads to the partial loss of immune tolerance.

## Introduction

Infection with Hepatitis B virus (HBV) causes a wide spectrum of disease manifestations, ranging from asymptomatic infection to acute self-limiting or fulminant hepatitis, or chronic infection with variable disease activity. Partially chronic infected patients may eventually develop cirrhosis or hepatocellular carcinoma. HBV is the prototype member of the hepadnavirus family. It is a small DNA virus with a 3.2-kb genome, which contains four genes named S, X, P and C genes. The S gene codes for the three co-carboxy-terminal envelope proteins termed large (LHBs), middle (MHBs) and small surface antigens (SHBs) or pre-S1, pre-S2 and major S proteins. The X gene codes for a 17-kDa regulatory protein. The P gene codes for the viral DNA polymerase, which is also a reverse transcriptase. The C gene codes for the core protein, which forms the viral core particle, and a related protein named the precore protein, which is the precursor of the secreted e antigen (HBeAg)[1]. HBeAg is found in serum during active infection and generally correlates with the degree of viremia. Indeed, HBV DNA level have been found to decline in serum following the development of an immuno response to HBeAg (anti-HBeAg)[2].

Previous studies indicated that Chronic HBV infection is associated with the emergence of mutations throughout the viral genome that result in the generation of diverse viral populations or quasispecies[3]. Point mutations unable to produce e antigen often become the dominant viral quasispecies in viral population present in the infected individual. The mechanism for their increased fitness may involve selection via the immune response or by enhancement of viral replication rates. The mutations associated with the loss of e antigen production in fulminant hepatitis patients frequently include the double mutation of A to T (A→T) at nt 1762 and G to A (G→A) at nt 1764 basal core promoter (BCP) mutation[4], and the G to A (G→A) at nt 1896 pre-C stop-codon mutation[5], the latter often accompanied by a G to A (G→A) mutation at nt 1899. Moreover, additional mutation in the BCP region that may confer increased replication efficiency to the virus have been described[6, 7].

It has been postulated that fulminant induced by HBV infection may be the result of increased viral replication results from the BCP mutation, which up-regulate pgRNA, with concurrent downregulation of pre-C mRNA synthesis[6, 8]. However, in vitro experiments addressing these issues have provided conflicting results[6, 9]. Some studies have reported increased viral replication in conjunction with a decrease in pre-C mRNA synthesis[6, 8], whilst others reported no increase in viral replication, only reduced pre-C mRNA production[10]. Such studies have employed reporter-gene constructs containing subgenomic fragments bearing the relevant mutations in the genetic background that they were found in or have used infectious clones with the mutations engineered by site-directed mutagenesis.

In the present study, BCP mutations and Pre-C mutations were separately introduced by site-directed mutagenesis in the same genetic background of an infectious clone, to determine the effect of these mutations per se on replication. In contrast to earlier studies, various infectious constructs were also introduced into mouse liver by hydrodynamic injection. Replication efficiency *in vivo* of various infectious constructs was measured by Southern Blot, Northern Blot, quantitative real-time PCR measurements of intracellular replicative intermediates, including RC dsDNA and pgRNA and released virion-associated HBV DNA. HBV DNA replication persistence was also detected in mouse.

## Materials and Methods

### Plasmid constructs and site-directed mutagenesis

A replication-competent plasmid, named pHBV1.2-WT, containing 1.2 copies genome length of HBV subtype adr (genotype C) was kindly provided by Peter Karayiannis[11]. This plasmid formed the template for use with a GeneArt^®^ Site-Directed Mutagenesis System (ThermoFisher, cat. #: A13282) according to the manufacturer’s instructions. A plasmid, named pHBV1.2-BCP, that contained an A to T mutation at nucleotide 1762 and G to A mutation at nucleotide 1764 was constructed. A plasmid, named pHBV1.2-PC, that contained a G to A mutation at nucleotide 1896 was constructed. A compensatory mutation of C to U at nucleotide 1858 was also created to maintain the stability of the overlapping epsilon structure[12].

### Cell culture and transfection

HepG2 cells were maintained in DMEM medium supplemented with 10% of fetal bovine serum, 100 μg/ml penicillin and 100 μg/ml streptomycin. Cells were seeded in 6-well plates at approximately 60% confluence. After 24 h, cells were transfected with Lipofectamine 2000 (Invitrogen, Camarillo, USA) according to the manufacturer’s instructions. The efficiency of transfection procedure was verified and controlled by co-transfection of a reporter plasmid pVIVO-Lucia/SEAP plasmids (Promega, USA), which expresses luciferase and SEAP.

### Hydrodynamic Injection of plasmids

Eight-week old male C57BL/6 mice were purchased from the experimental animal center in Henan Province, China. Our animal protocol was approved by the Life Science Ethics Committee of Zhengzhou University. Mice were injected via the tail vein with plasmids in 5-8 seconds in a volume of saline equivalent to 8% of the body weight of the mouse as described previously[13]. pVIVO-Lucia/SEAP plasmids (invivoGen, San Diego, CA, USA), which expresses the luciferase and SEAP, was included in the injection to serve as the internal control to monitor the injection efficiency.

### Southern and Northern blot analyses

Transfected cells or Liver tissues were homogenized in DNA lysis buffer (20 mM Tris-HCl, pH7.0, 20 mM EDTA, 50 mM NaCl, 0.5% SDS), incubated for 16 hours at 37°C with proteinase K (600 μg/ml) and then phenol/chloroform extracted for the isolation of DNA. The HBV RI DNA in the core particles was isolated using previous protocol [28]. For RNA isolation, liver tissues were homogenized in Trizol (Invitrogen) and total RNA was isolated following the manufacturer’s protocol. Both Southern and Northern blot analyses were conducted using the ^32^P-labeled HBV DNA probe.

### Real-time PCR analysis of medium or serum HBV DNA

100 μl cell culture medium or 10 μl mouse serum was digested with 10 μg DNase I and micrococcal nuclease for 30 min at 37°C to remove free DNA. Then added 100 μl lysis buffer (20 mM Tris-HCl, 20 mM EDTA, 50 mM NaCl, and 0.5%SDS) containing 27μg proteinase K. After incubation at 65°C overnight, viral DNA was isolated by phenol/chloroform extraction and ethanol precipitation. The DNA pellet was rinsed with 70% ethanol and resuspended in 10 μl TE (10 mM Tris-HCl [pH 7.0], 1 mM EDTA). HBV DNA was then isolated as described above. For HBV real-time PCR analysis, the following primers were used: forward primer, 1552-CCGTCTGTGCCTTCTCATCTG-1572; and reverse primer, 1667-AGTCCTCTTATGTAAGACCTT-1646. The TaqMan probe used was 1578-CCGTGTGCACTTCGCTTCACCTCTGC-1603.

### HBV serological assays by ELISA

HBsAg and HBeAg were analyzed using the ELISA kit following the manufacturer’s instructions (Kehua, China). The mouse serum was diluted 50-fold with PBS, and 100 μl of diluted sample was used for the assay. The assays were conducted in triplicate.

### Isolation of mononuclear cells from the mouse liver

The mouse liver was perfused with 30 ml PBS via the hepatic portal vein at room temperature. The liver was excised, and liver cells were dispersed in PBS containing 2% fetal bovine serum (FBS) and 0.02% NaN_3_ at 4°C through a 40-μm nylon cell strainer (DN Falcon #352340). Cells were centrifuged at 500xg at 4°C for 5 minutes. Liver mononuclear cells were then isolated by centrifugation at 700×g in 25 ml of 33.75% sterile Percoll at 25°C for 7 minutes. Cells were incubated with the 1× Red Blood Cell lysis buffer (Sigma) at room temperature for 4 minutes, and washed three times with PBS containing 2% FBS and 0.02% NaN_3_ at 4°C with centrifugation at 500×g for 5 minutes after each wash. After the first wash, cells were again filtered through a 40-μm nylon cell strainer and, after the last wash, cells were resuspended in 1 ml PBS containing 2% FBS and 0.02% NaN_3_ and counted.

### Flow cytometry

The following antibodies, which were all purchased from eBioscience, San Diego, were used for flow cytometry: anti-mouse CD3 eFluor 450 (cat. 48-0032-82), anti-mouse CD4 FITC (cat. 11-0042-82), anti-mouse CD25 PE (12-0251-82), and anti-mouse FoxP3 APC (cat.17-5773-82). Two million cells per sample were used for flow cytometry. Cells were incubated with Fc Block for 15 minutes at 4°C in darkness to block the FcγR and then stained with fluorochrome-conjugated antibodies for 30 minutes at 4°C also in darkness. Cells were then analyzed using a Canto II (BD Bioscience, CA) 8-color flow cytometer, and data were analyzed using FlowJo software (Tree Star, Inc., Ashland, OR).

### The luciferase reporter assay

The reporter construct pVIVO-Lucia/SEAP was transfected into HepG2 cells or delivered into the mouse liver by hydrodynamic injection. The cells culture medium was diluted 100 folds; the mouse serum was diluted 5000 folds. SEAP expression was be quantified by using The SEAP Reporter Assay Kit (invivoGen, San Diego, CA, USA). All the experiments were repeated at least three times.

### Statistical Analyses

Data were presented as the mean ± SEM, and *p* values were determined by two-tailed Student’s *t* test using GraphPad Prism software. Differences that were statistically significant were indicated by asterisks in the figures.

## Results

### Replication capacity of BCP and pre-C constructs in vitro

The construction pHBV1.2-BCP contains double mutation of A to T (A→T) at nt 1762 and G to A (G→A) at nt 1764 basal core promoter (BCP) mutation, and the construction pHBV1.2-PC contain G to A (G→A) at nt 1896 pre-C stop-codon mutation were introduced by site-directed mutagenesis in the same genetic background of a HBV replication-competent plasmid pHBV1.2-WT, the latter accompanied by a G to A (G→A) mutation at nt 1899. To examine the replication capacity of each construction, each construction was transfected into HepG2 cells. Replication capacity of each construct was analyzed in terms of HBV core particle DNA and total cellular RNA by Southern and Northern Blot in transfected HepG2 cells at 72 hours post-transfection. As shown in figure 1A, the level of HBV core particle DNA was increased by 40% in construct with the BCP mutations, increased 1.8 folds in construct with Pre-C mutation comparing with that in pHBV1.2-WT transfected cells. When the HBV RNA was analyzed by Northern blot, a significant increase of the HBV RNA level both in BCP mutations and Pre-C mutation was also observed, particularly with the HBV S gene transcripts. The HBsAg and HBeAg in the supernatant of cell culture were also measured by ELISA. The results shown these mutations did not affect the expression of HBsAg in HepG2 cells, but Pre-C mutation abolished the expression of HBeAg, and BCP double mutations reduce HBeAg exprsssion about 50% (Figure 1B). The viral titer in the supernant of cell culture transfected BCP or stop-codon mutations inceased about 3 tines and 7 times than that of in cell supernant transfected with wildtype replicated HBV DNA (Fig. 1C)

**Figure 1.**
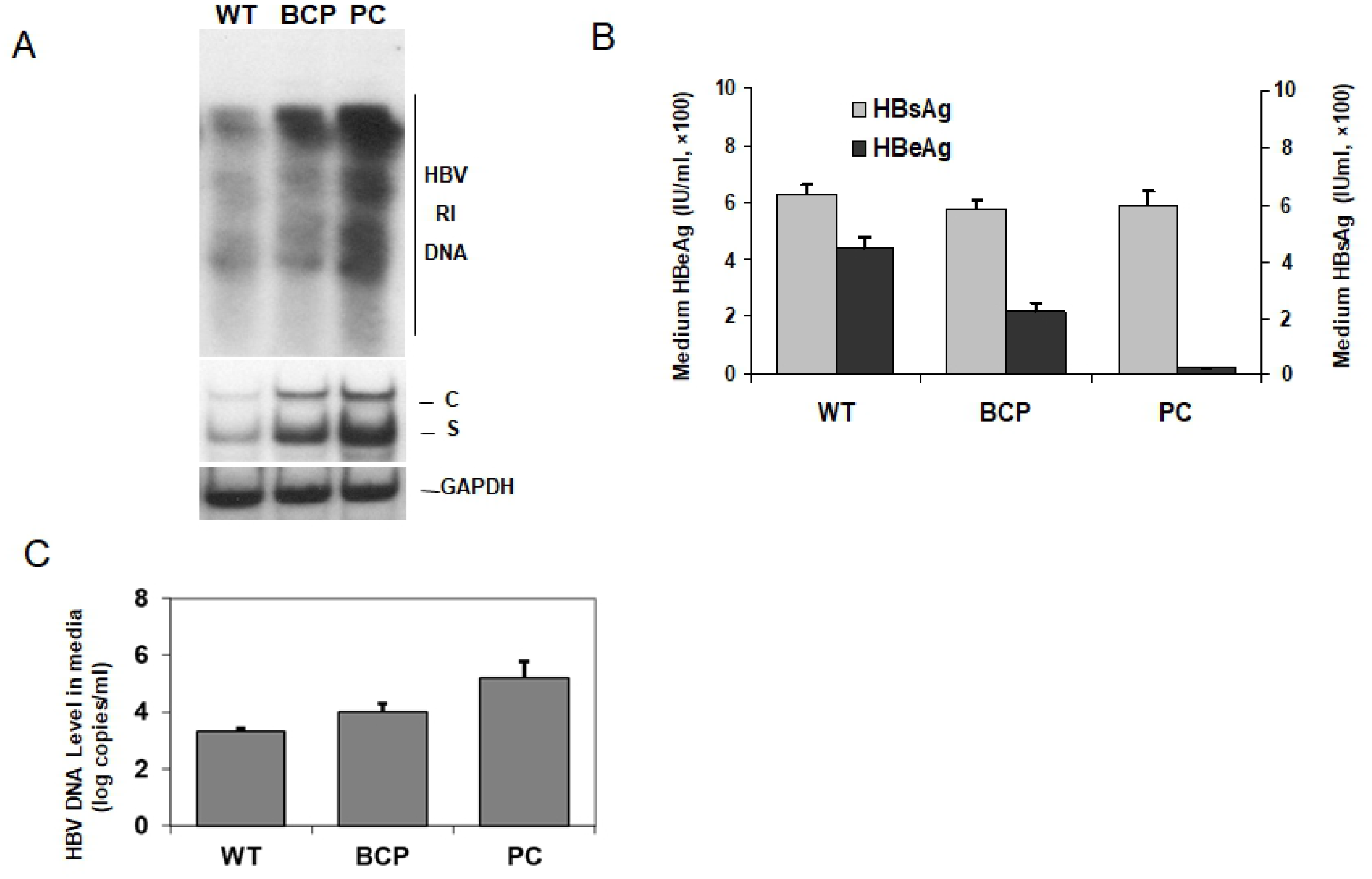
The effect of BCP or Pre-C mutations on HBeAg expression and HBV replication in vitro. HepG2 cells were transfected with the pHBV1.2-WT, pHBV1.2-BCP or pHBV1.2-PC for 48 hours. (A) The cells were lysed for the analysis of the HBV RI DNA by Southern-blot and HBV RNA by Northern Blot. (B) The incubation media were then harvested for the analysis of HBsAg and HBeAg by ELISA. (C) HBV DNA levels in the media was analyzed by real-time PCR.

### Replication capacity of BCP and pre-C constructs in vivo

To examine the replication capacity of each construction in vivo, we performed the hydrodynamic injection in mice, which is a rapid and convenient method for HBV gene delivery into mouse liver. 4 μg pHBV1.2-WT, pHBV1.2-BCP or pHBV1.2-PC plasmids were injected via the tail vein into the mice, respectively. HBsAg, HBeAg and the viral titer in the serum were measured by ELISA and real-time PCR at 72 hours after injection. As shown in Figure 2A, HBsAg level increased about 1 time in the mice injected with pHBV1.2-BCP plasmid, and 1.5 times in the mice injected with pHBV1.2-PC plasmid compared with that in mice injected with pHBV1.2-WT control HBV DNA. HBeAg level decreased about 2 time in the mice injected with pHBV1.2-BCP plasmid compared with that in mice injected with control plasmid, and HBeAg was undectable in the mice injected with pHBV1.2-PC plasmid. HBV titer in the mice serum was also detected by qPCR. As shown in Figure 2B, HBV titer increased in mice injected with either pHBV1.2-BCP or pHBV1.2-PC plasmid compared with that in mice injected with pHBV1.2-WT, but increased more significantly in the mice injected with pHBV1.2-PC.

**Figure 2.**
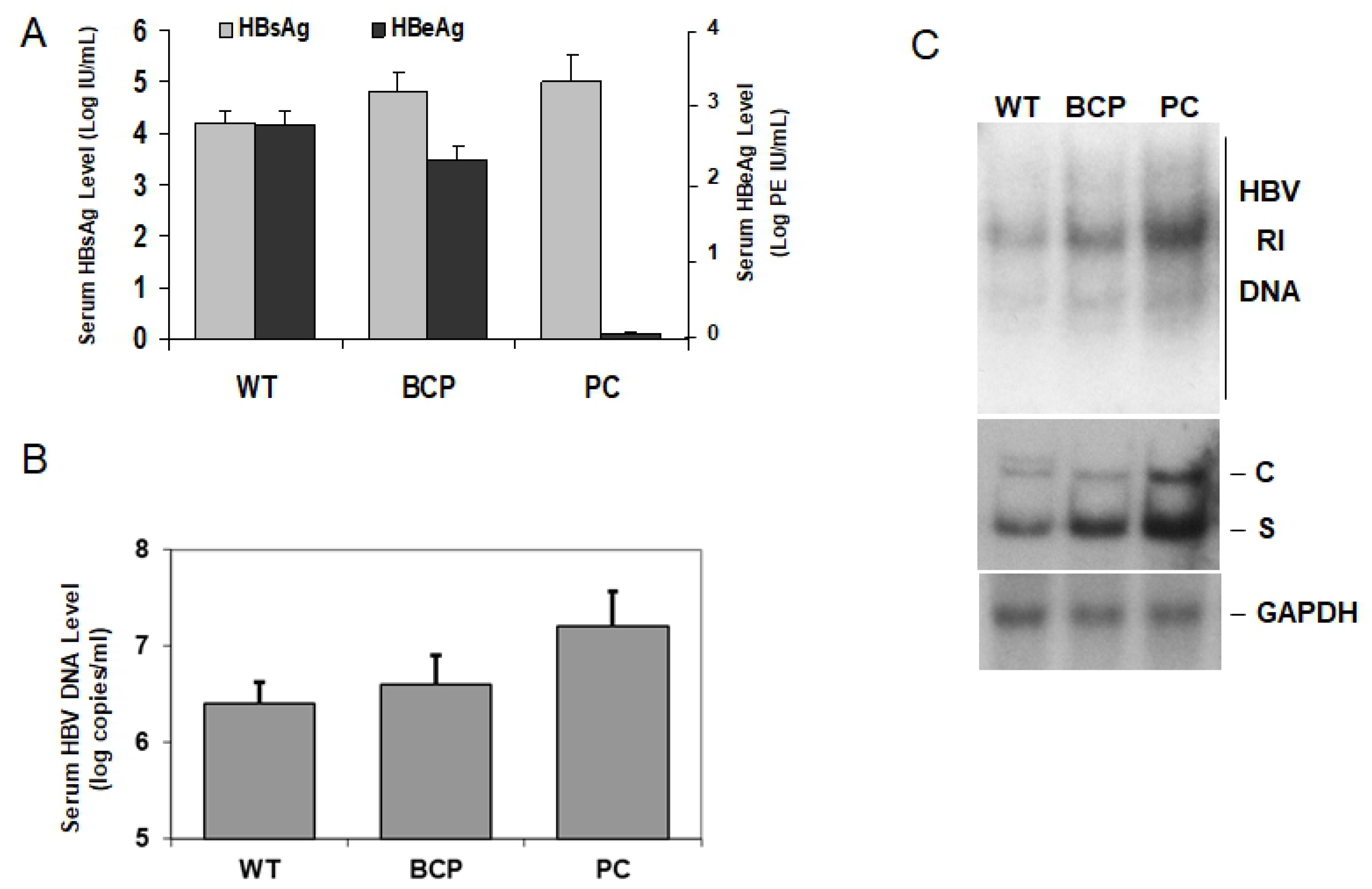
The effect of BCP or Pre-C mutations on HBeAg expression and HBV replication in vivo. 4 μg pHBV1.2-WT, pHBV1.2-BCP or pHBV1.2-PC plasmids were injected via the tail vein into the mice by hydrodynamic injection, respectively. Mice sacrificed at 72 hours after injection. (A) HBsAg and HBeAg in the serum were measured by ELISA. (B) HBV DNA levels in the serum was analyzed by real-time PCR. (C) 50 ug the liver tissues were lysed for the analysis of the HBV RI DNA by Southern-blot or HBV RNA by Northern Blot.

HBV core particle DNA and total cellular HBV RNA were then analyzed. As showin in the figure 2C, both BCP and Pre-C mutations led to an increasing level of HBV core partical DNA and HBV RNA, but PC mutations had a more prominent effect on HBV core partical DNA and HBV RNA.

### BCP and Pre-C mutations were not benefit for HBV persistence in mouse

Previous studies demonstrated that HBeAg may be important for HBV persistence after vertical transmission [2, 14, 15]. Expression of HBeAg was also important for HBV to establish persistent replication in mice. As BCP mutants partialy reduce HBeAg level and Pre-C mutants abolish HBeAg expression, We want to know whether BCP and Pre-C mutants affect HBV persistence replication in mice. 4 μg pHBV1.2-WT, pHBV1.2-BCP or pHBV1.2-PC plasmids were injected via the tail vein into the mice, respectively. Mouse blood was collected at different time point after injection. HBsAg and the viral titer in the serum were measured by ELISA and real-time PCR, respectively. As shown in Figure 3A, mice injected with pHBV1.2-WT, pHBV1.2-BCP or pHBV1.2-PC plasmids produced initially a high level of HBsAg in the serum. This HBsAg level declined rapidly in mice injected with pHBV1.2-BCP or pHBV1.2-PC plasmids. HBsAg became undetectable in three weeks in the mice injected with pHBV1.2-PC plasmid. HBsAg became undetectable in five weeks in the mice injected with pHBV1.2-BCP plasmid. In contrast, the HBsAg level in mice injected with pHBV1.2-WT plasmid declined slowly and was still detectable at 7 weeks after injection, the study endpoint. The level of virion-associated HBV DNA in the serum peaked at 1 day post-infection in mice injected with pHBV1.2-WT, pHBV1.2-BCP or pHBV1.2-PC plasmids. It was approixmately 6.3×10^6^ genome copies/ml. HBV DNA in the serum declined rapidly and became undetectable in mice injected with pHBV1.2-PC mutant plasmid after 3 weeks. In mice injected with pHBV1.2-BCP plasmid, the virion-associated HBV DNA became undetectable after 5 weeeks. However, in the mice imjected with pHBV1.2-WT, the virion-associated HBV DNA reduced slightly after day 4 but remained more or less at the same level at about 10^3^ genome copies/ml up to 7 weeks after injectionim. Mouse liver HBV DNA and RNA was also confirmed by detecting HBV serun DNA replicative intermediates (RI). As shown in figure 3B, the HBV RI DNA could be detected by Southern blot in the mouse liver at 6 weeks post-injection of pHBV1.2. In contrast, HBV RI DNA could not be detected in the mouse liver injected with pHBV1.2-BCP or pHBV1.2-PC at same time point. Similarly, the HBV core protein could be detected by immunostaining in a significant number of hepatocytes (Figure 3C) in mouse at 6 weeks after pHBV1.2-WT injection. In contrast, the HBV core protein could not be detected in the hepatocytes of mice injected with pHBV1.2-BCP or pHBV1.2-PC at the same time point after injection. These results that HBV could not persistently replicate longer in mice injected with pHBV1.2-BCP or pHBV1.2-PC than that in mice injected with pHBV1.2 indicated an important effect of HBeAg on the establishment of HBV persistence in mice.

**Figure 3.**
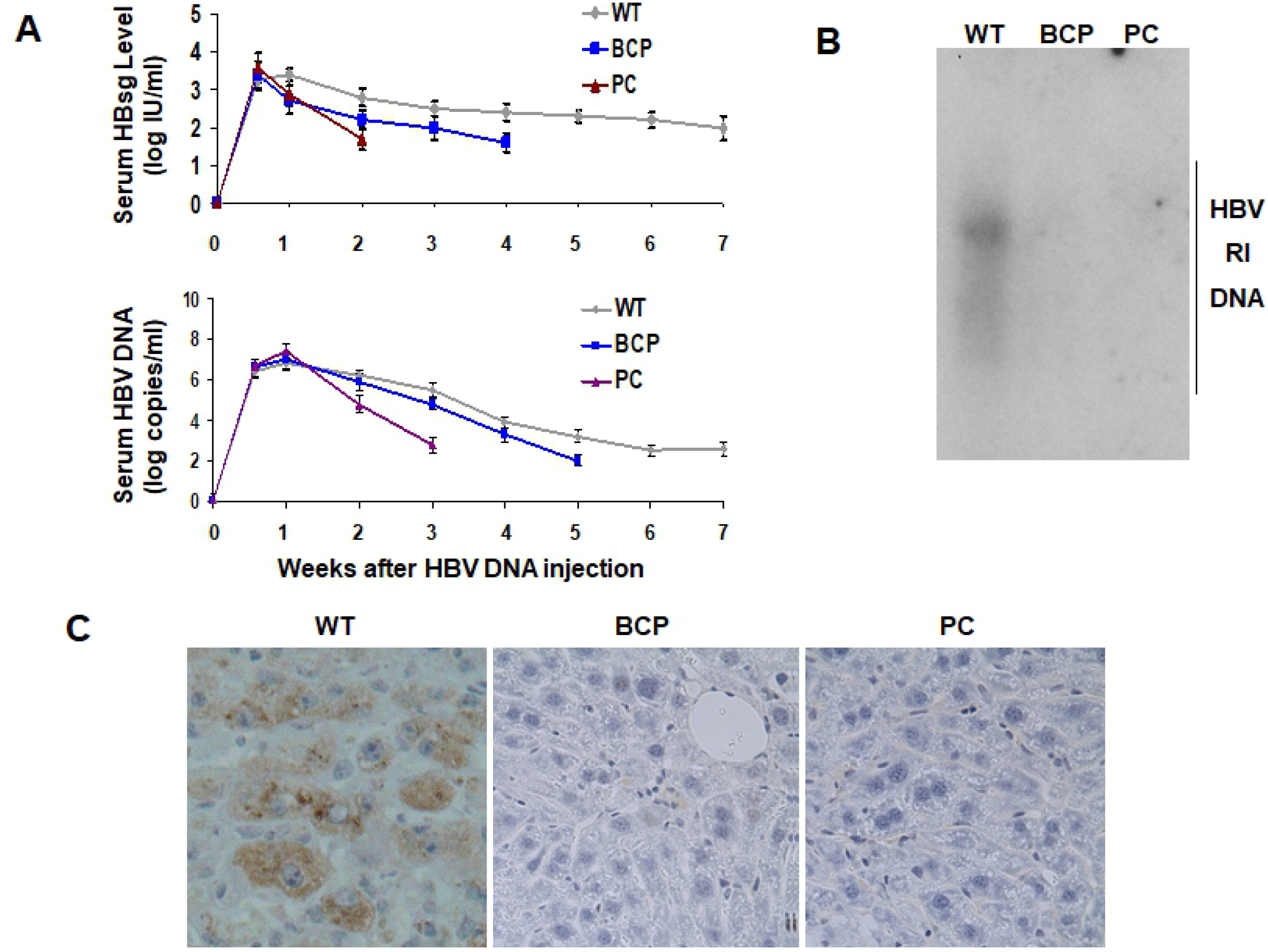
The effect of BCP or Pre-C mutations on HBV Persists in Mice. (A) Analysis of HBV persistence in pHBV1.2-WT, pHBV1.2-BCP or pHBV1.2-PC plasmids injected mice. The sera of 15 pHBV1.2-WT injected mice, 13 pHBV1.2-BCP injected mice and 16 pHBV1.2-PC injected mice were collected at the time points indicated and analyzed for HBsAg by ELISA (upper panel) and HBV DNA by real-time PCR (lower panel). (B) HBV replicative intermediate (RI) DNA in the mice liver was analyzed by Southern blot. The liver tissues of mice were isolated at 6 weeks after HBV DNA injection for the analysis. (C) Immunostaining of HBV core protein in the liver of different mice. The liver tissues of mice were collected at 6 weeks after HBV DNA injection for the analysis.

### Pre-C mutations increased CD8+ CTL response in mice

To understand why HBV persisted shorter in mice injected with pHBV1.2-PC than that in mice injected with pHBV1.2-WT or pHBV1.2-BCP plasmids, we firstly analyzed the level of HBV-associate CD8+ T cells in mice two weeks after the injection of HBV DNA. Mononuclear cells isolated from the liver were stimulated with a peptide derived from the HBV core protein (amino acids 93–100, MGLKIRQL) as previously described [13]. The IFN-γ-producing CD8+ T cells were then analyzed by flow cytometry. As show in Figure 4A, control mice injected with the pUC19 vector DNA had few activated interferon-γ (IFN-γ)-CD8+ T cells stimulating with the HBV peptide. For the mice injected with the pHBV1.2, the population of IFN-γ-positive CD8+ T cells was 1.7% in the liver. The population of IFN-γ-positive CD8+ T cells was 1.9% in the liver of mice injected with pHBV1.2-BCP. But the population of IFN-γ-positive CD8+ T cells was 3.2% in the liver of mice injected with PC mutant HBV DNA.

**Figure 4.**
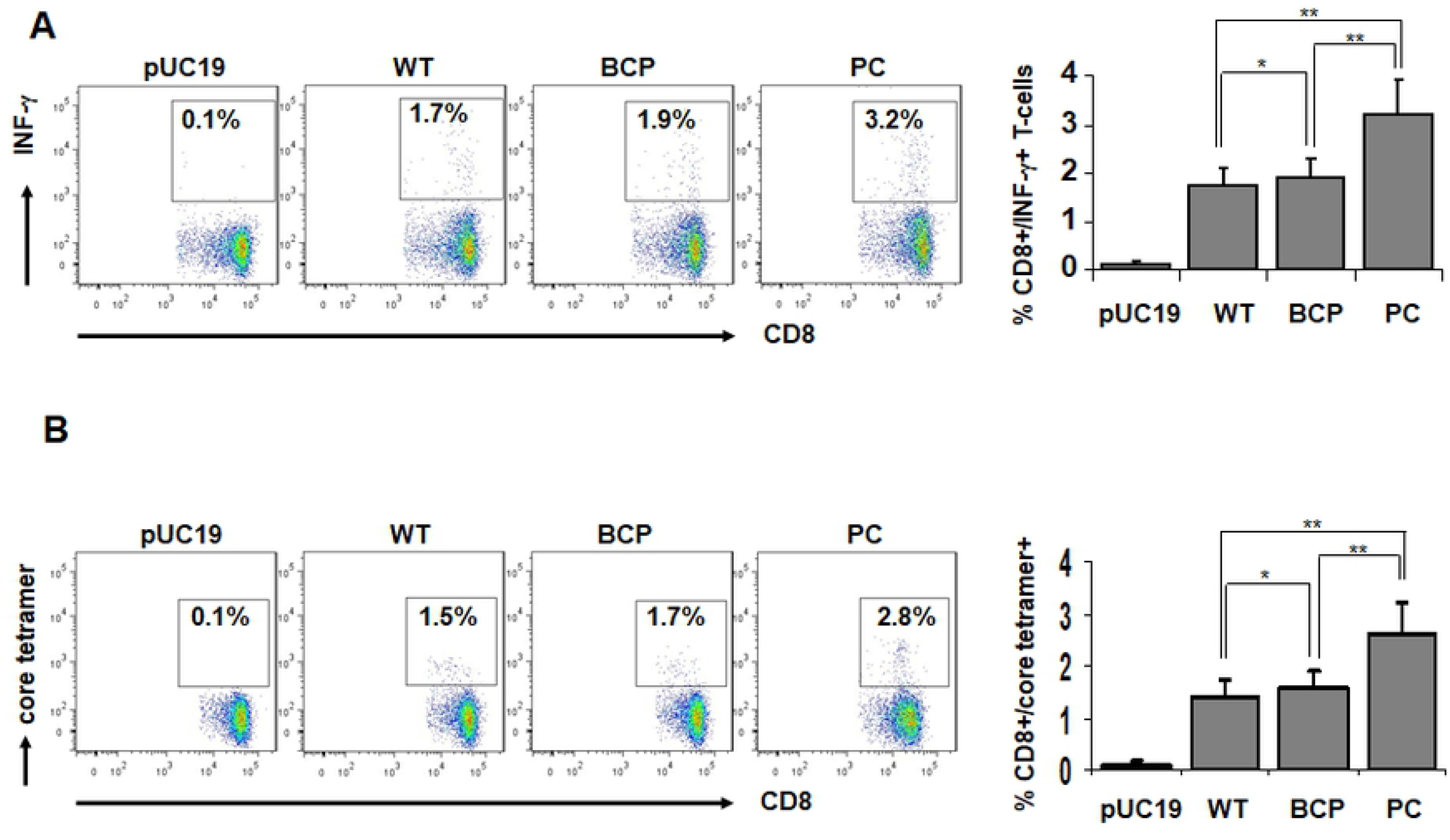
The effect of BCP or Pre-C mutations on CTL responses in mice. (A) Analysis of HBV-specific CD8^+^ T cells. 4 μg pUC19, pHBV1.2-WT, pHBV1.2-BCP or pHBV1.2-PC plasmids were injected via the tail vein into the mice by hydrodynamic injection, respectively. Mice were sacrificed at 14 days after injection for the isolation of liver mononuclear cells. These cells were then stimulated with the HBV core peptide (aa 93-100) and analyzed by flow cytometry for CD8^+^IFN-γ^+^ cells. The results represent the mean values of five different mice. (B) Total number of HBV-specific CD8+ T cells per mouse liver. CD8^+^ T cells stained by the HBV core tetramer were analyzed by flow cytometry. The results represent the mean of five different mice. *p > 0.05; **p < 0.01.

We also analyzed the HBV-specific hepatic CD8+ T cells with the tetramer containing a peptide derived from the HBV core protein as described above. The results revealed a slightly increase of HBV-specific CD8+ T cells in mice injected with pHBV1.2-PC comparing with the mice injected with pHBV1.2. As show in Figure 4B, in mice injected with the pHBV1.2-WT or pHBV1.2-BCP plasmid, the populations of HBV-specific CD8+ T cells in the liver were 1.5% and 1.7%, respectively. In contrast, a significant increase of the population of HBV-specific CD8+ T cells was observed in the liver of mice injected with pHBV1.2-PC plasmid (2.8%). As Pre-C mutation abolished HBeAg express, these results suggested that the increased CD8+ CTL response in pHBV1.2-PC injected mice might be due to the lack of HBeAg in those cells.

### Pre-C mutations decreased the population of Regulatory T cells (Tregs) in mice

Tregs have an important role in regulating effector T cell responses in many infectious diseases[16, 17]. Tregs have been reported to down-regulating HBV-specific T cell responses in HBV infected patients as well as in HBV animal models[18, 19]. To understand whether stronger HBV-specific T cell responses in mice injected with pHBV1.2-PC was related to Tregs. Treg response in the spleen and liver were analyzed by Flow cytometry. 14 days after mice hydrodynamic injection, mice liver monocytes and spleen cells were isolated and absolute numbers of CD4+ Foxp3+ Treg cells were analyzed by flow cytometry. As show in Figure 5, injection of pHBV1.2-WT, pHBV1.2-BCP or pHBV1.2-PC plasmids resulted in a slightly increasing population of CD4+ Foxp3+ Treg+ cells in mice spleen comparing with that in control mice injected with pUC19 plasmid. The population of Treg cells in liver significantly increased in the mice injected with pHBV1.2-WT comparing with that in mice injected with pUC19, pHBV1.2-BCP or pHBV1.2-PC. The population of Treg cells in liver slightly increased in the mice injected with pHBV1.2-BCP or pHBV1.2-PC comparing with that in mice injected with pUC19. As Pre-C mutation abolished the expression of HBeAg and BCP mutations reduce HBeAg exprsssion both in vitro or in vivo, these results demostrated that HBeAg might has an important role in HBV persistence by regulating Treg popula

**Figure 5.**
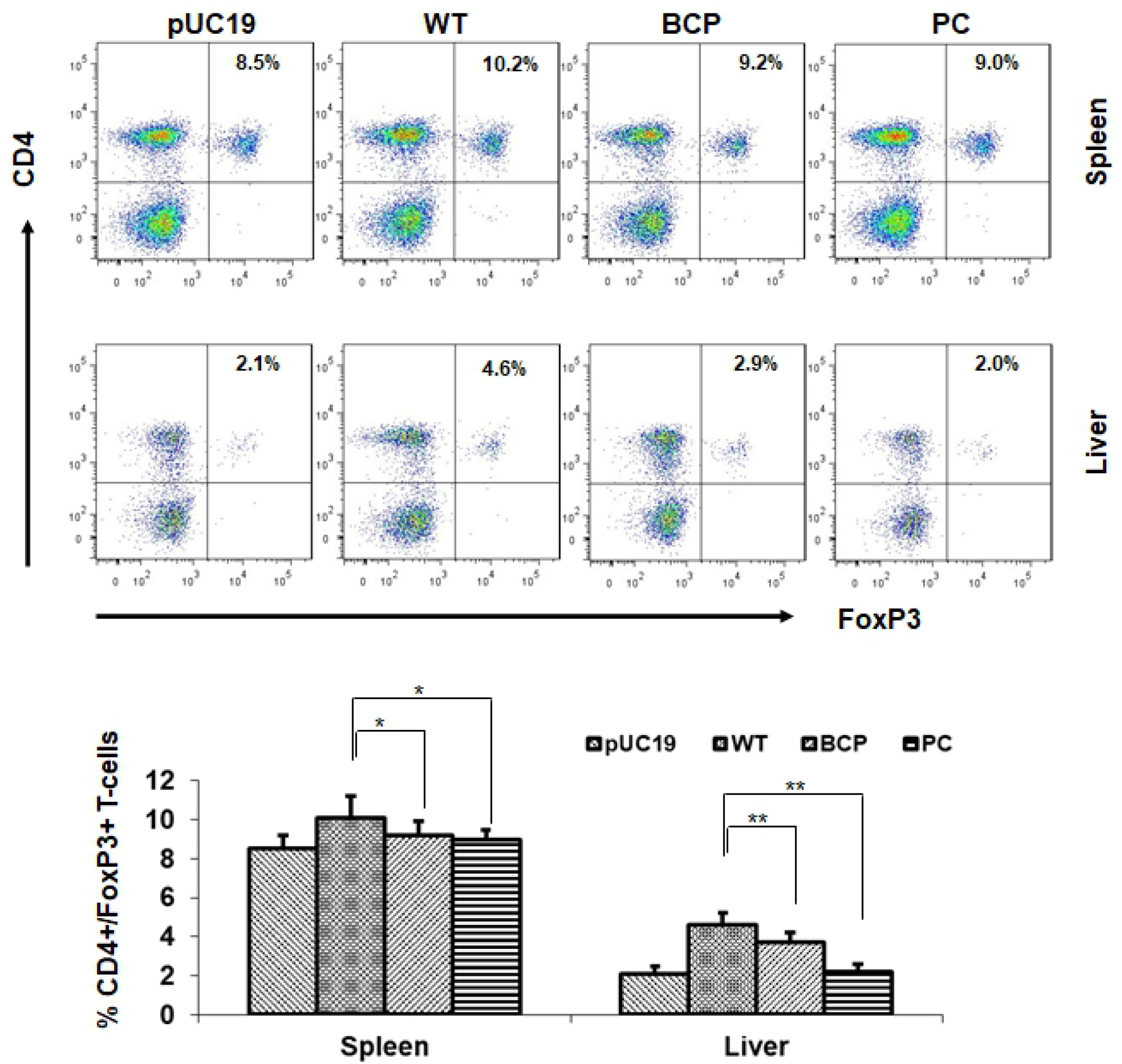
The effect of BCP or Pre-C mutations on Treg in mice. 4 μg pUC19, pHBV1.2-WT, pHBV1.2-BCP or pHBV1.2-PC plasmids were injected via the tail vein into the mice by hydrodynamic injection, respectively. Mice were sacrificed at 14 days after injection for the isolation of liver mononuclear cells and spleen cells. CD3^+^/CD4^+^/FoxP3^+^ Treg were analyzed by flow cytometry. The results represent the mean of five different mice. *p > 0.05; **p < 0.01.

## Discussion

The appearance of the BCP or Pre-C mutations, which reduce or abolish HBeAg production[9, 20], heralds the initiation of the seroconversion phase from HBeAg to anti-HBeAg positivity in clinical[21]. The remove of the HBeAg often lead to the awakening of the immune response[22]. However, the appearance of the BCP or Pre-C mutations also often result to HBeAg negative chronic hepatitis B with high viremia levels[23]. Many studies have confirmed that BCP mutations reduce HBeAg expression and pre-C mutations abrogate HBeAg synthesis in vitro[11, 24]. However, there is still no clear evidence to show whether the reduction or loss of HBeAg caused by BCP or Pre-C mutations is associated with HBV persistence. In this study, we constructed a replication-competent HBV plasmid containing BCP and Pre-C mutations, respectively. The two plasmids were transfected into HepG2 cells. The BCP mutation reduced the HBeAg level by 50% and increased the HBV replication level about 3 times. The Pre-C mutation completely abolished HBeAg expression, and the HBV replication level increased 7 times compared with the control. These results consist with previous results[11, 24, 25]. To further study the effect of BCP and Pre-C mutations on HBV replication in vivo, we injected these two mutant plasmids into the liver of mice by hydrodynamic injection. The results showed that the BCP mutation reduced the serum HBeAg level, and the Pre-C mutation eliminated HBeAg expression in mice. The BCP mutation slightly increased the levels of HBV DNA and RNA in the liver of mice, and the HBV titer in the serum. The Pre-C mutation significantly increased both HBV DNA and RNA levels in the liver of mice, and the level of HBV DNA in the serum increased more significantly.

As the BCP and Pre-C are frequently observed in HBV sequences isolated from chronic patients[4, 5], we want to know whether the BCP and Pre-C mutations contribute to HBV persistent infection. In this study, we injected low density (4μg/mouse) pHBV1.2-WT, pHBV1.2-BCP or pHBV1.2-PC into mice, respectively. Our results shown that HBV could persist in mice injected with pHBV1.2-WT for about 7 weeks. However, HBV could persist about 5 weeks in mice injected with pHBV1.2-BCP, and 3 weeks only in mice injected with pHBV1.2-PC plasmid. As BCP and Pre-C mutations affect HBV replication level and the expression of HBeAg, which has a role of regulating immune activity, we believe that the reduction or absence of HBeAg induced by BCP or Pre-C mutations significantly shortens the HBV persistence in mice.

Previous studies shown that HBeAg plays an important role in the development of CHB by regulating the host immune response through multiple pathways. Circulating HBeAg may downregulate antiviral clearance mechanisms by virtue of eliciting anti-inflammatory Th2-like cytokine[26] and deplete inflammatory HBeAg-specific Th1 cells[27]. In addition, studies with mouse models have shown that HBeAg impairs liver HBV-specific CD8^+^ cytotoxic T lymphocytes mediated by hepatic macrophages[28].These results indicated that HBeAg has the importance role in HBV persistence. In current study, we analyzed the immune responses in pUC19, pHBV1.2-WT, pHBV1.2-BCP or pHBV1.2-PC plasmids injected mice. We found the HBV related CD8+ CTL response in pHBV1.2-BCP injected mice only slightly increased comparing with that in mice injected with pHBV1.2. However, the HBV related CD8+ CTL response in pHBV1.2-PC injected mice significant increased compared with that in control mice. These results indicated that loss of HBeAg might upregulate HBV related CD8+ CTL response which terminate HBV persistent.

To understand why HBV specific CD8+ CTL response in pHBV1.2-PC injected mice is stronger than that in pHBV1.2-WT injected mice, we analyzed the CD3^+^CD4^+^Foxp3^+^ regulatory T cells (Tregs), as several reports have suggested that it plays a significant role in suppressing T cell responses during viral infections[16, 17]. Some studies also indicated that Tregs play an important role in down-regulating HBV-specific effector T cell responses in HBV patients[18, 19, 29]. In addition, studies with mouse models have shown that absolute Treg numbers significantly increased in the liver after hydrodynamic injection of replicative HBV DNA[30]. In this study, we found that population of Tregs significantly increased in liver but not in spleen of mice injected with pHBV1.2-PC comparing with that in mice injected with pHBV1.2-WT or pHBV1.2-BCP. It indicates that HBeAg might contribute to regulation of Treg population only in liver, which affect HBV specific CD8+ CTL response and HBV persistence.

Taken together, in this study we demonstrated that HBeAg play an important role in suppressing the CTL response, which is related with increasing the frequency of Tregs in mouse. Lack of HBeAg expression leads to the partial loss of immune tolerance. This study will provide important information for us to understand the mechanism of HBV persistence and to improve the treatments for chronic HBV patients.

## Author Contributions

C.Y and L. JP designed, performed study, Z. WX and Y. LZ analyzed the date. Y. K and T. YJ directed the study and wrote the manuscript.

## Acknowledgements

We wish to thank all the subjects volunteered to participate in this study. This work was supported by the National Natural Science Foundation of China (U1704179).

## References

1. Ou JH, Laub O, Rutter WJ. Hepatitis B virus gene function: the precore region targets the core antigen to cellular membranes and causes the secretion of the e antigen. Proc Natl Acad Sci U S A. 1986;83(6):1578–82. Epub 1986/03/01. doi:10.1073/pnas.83.6.1578. PubMed PMID:3006057; PubMed Central PMCID:PMCPMC323126.

2. Milich D, Liang TJ. Exploring the biological basis of hepatitis B e antigen in hepatitis B virus infection. Hepatology. 2003;38(5):1075–86. Epub 2003/10/28. doi:10.1053/jhep.2003.50453. PubMed PMID:14578844.

3. Revill PA, Tu T, Netter HJ, Yuen LKW, Locarnini SA, Littlejohn M. The evolution and clinical impact of hepatitis B virus genome diversity. Nat Rev Gastroenterol Hepatol. 2020. Epub 2020/05/30. doi:10.1038/s41575-020-0296-6. PubMed PMID:32467580.

4. Okamoto H, Tsuda F, Akahane Y, Sugai Y, Yoshiba M, Moriyama K, et al. Hepatitis B virus with mutations in the core promoter for an e antigen-negative phenotype in carriers with antibody to e antigen. J Virol. 1994;68(12):8102–10. Epub 1994/12/01. doi:10.1128/JVI.68.12.8102-8110.1994. PubMed PMID:7966600; PubMed Central PMCID:PMCPMC237274.

5. Carman WF, Jacyna MR, Hadziyannis S, Karayiannis P, McGarvey MJ, Makris A, et al. Mutation preventing formation of hepatitis B e antigen in patients with chronic hepatitis B infection. Lancet. 1989;2(8663):588–91. Epub 1989/09/09. doi:10.1016/s0140-6736(89)90713-7. PubMed PMID:2570285.

6. Baumert TF, Marrone A, Vergalla J, Liang TJ. Naturally occurring mutations define a novel function of the hepatitis B virus core promoter in core protein expression. J Virol. 1998;72(8):6785–95. Epub 1998/07/11. doi:10.1128/JVI.72.8.6785-6795.1998. PubMed PMID:9658127; PubMed Central PMCID:PMCPMC109887.

7. Parekh S, Zoulim F, Ahn SH, Tsai A, Li J, Kawai S, et al. Genome replication, virion secretion, and e antigen expression of naturally occurring hepatitis B virus core promoter mutants. J Virol. 2003;77(12):6601–12. Epub 2003/05/28. doi:10.1128/jvi.77.12.6601-6612.2003. PubMed PMID:12767980; PubMed Central PMCID:PMCPMC156182.

8. Buckwold VE, Xu Z, Chen M, Yen TS, Ou JH. Effects of a naturally occurring mutation in the hepatitis B virus basal core promoter on precore gene expression and viral replication. J Virol. 1996;70(9):5845–51. Epub 1996/09/01. doi:10.1128/JVI.70.9.5845-5851.1996. PubMed PMID:8709203; PubMed Central PMCID:PMCPMC190601.

9. Li J, Buckwold VE, Hon MW, Ou JH. Mechanism of suppression of hepatitis B virus precore RNA transcription by a frequent double mutation. J Virol. 1999;73(2):1239–44. Epub 1999/01/09. doi:10.1128/JVI.73.2.1239-1244.1999. PubMed PMID:9882327; PubMed Central PMCID:PMCPMC103946.

10. Sterneck M, Kalinina T, Gunther S, Fischer L, Santantonio T, Greten H, et al. Functional analysis of HBV genomes from patients with fulminant hepatitis. Hepatology. 1998;28(5):1390–7. Epub 1998/10/31. doi:10.1002/hep.510280530. PubMed PMID:9794926.

11. Jammeh S, Tavner F, Watson R, Thomas HC, Karayiannis P. Effect of basal core promoter and pre-core mutations on hepatitis B virus replication. J Gen Virol. 2008;89(Pt 4):901–9. Epub 2008/03/18. doi:10.1099/vir.0.83468-0. PubMed PMID:18343830.

12. Lok AS, Akarca U, Greene S. Mutations in the pre-core region of hepatitis B virus serve to enhance the stability of the secondary structure of the pre-genome encapsidation signal. Proc Natl Acad Sci U S A. 1994;91(9):4077–81. Epub 1994/04/26. doi:10.1073/pnas.91.9.4077. PubMed PMID:8171038; PubMed Central PMCID:PMCPMC43726.

13. Yang PL, Althage A, Chung J, Maier H, Wieland S, Isogawa M, et al. Immune effectors required for hepatitis B virus clearance. Proc Natl Acad Sci U S A. 2010;107(2):798–802. Epub 2010/01/19. doi:10.1073/pnas.0913498107. PubMed PMID:20080755; PubMed Central PMCID:PMCPMC2818933.

14. Okada K, Kamiyama I, Inomata M, Imai M, Miyakawa Y. e antigen and anti-e in the serum of asymptomatic carrier mothers as indicators of positive and negative transmission of hepatitis B virus to their infants. The New England journal of medicine. 1976;294(14):746–9. doi:10.1056/NEJM197604012941402. PubMed PMID:943694.

15. Ou JH. Molecular biology of hepatitis B virus e antigen. Journal of gastroenterology and hepatology. 1997;12(9–10):S178–87. PubMed PMID:9407336.

16. Manigold T, Racanelli V. T-cell regulation by CD4 regulatory T cells during hepatitis B and C virus infections: facts and controversies. Lancet Infect Dis. 2007;7(12):804–13. Epub 2007/11/30. doi:10.1016/S1473-3099(07)70289-X. PubMed PMID:18045563.

17. Rowe JH, Ertelt JM, Way SS. Foxp3(+) regulatory T cells, immune stimulation and host defence against infection. Immunology. 2012;136(1):1–10. Epub 2012/01/04. doi:10.1111/j.1365-2567.2011.03551.x. PubMed PMID:22211994; PubMed Central PMCID:PMCPMC3372751.

18. Stoop JN, van der Molen RG, Baan CC, van der Laan LJ, Kuipers EJ, Kusters JG, et al. Regulatory T cells contribute to the impaired immune response in patients with chronic hepatitis B virus infection. Hepatology. 2005;41(4):771–8. Epub 2005/03/26. doi:10.1002/hep.20649. PubMed PMID:15791617.

19. Franzese O, Kennedy PT, Gehring AJ, Gotto J, Williams R, Maini MK, et al. Modulation of the CD8+-T-cell response by CD4+ CD25+ regulatory T cells in patients with hepatitis B virus infection. J Virol. 2005;79(6):3322–8. Epub 2005/02/26. doi:10.1128/JVI.79.6.3322-3328.2005. PubMed PMID:15731226; PubMed Central PMCID:PMCPMC1075696.

20. Koumbi L, Pollicino T, Raimondo G, Stampoulis D, Khakoo S, Karayiannis P. Hepatitis B virus basal core promoter mutations show lower replication fitness associated with cccDNA acetylation status. Virus Res. 2016;220:150–60. Epub 2016/05/02. doi:10.1016/j.virusres.2016.04.022. PubMed PMID:27132039.

21. Kamijo N, Matsumoto A, Umemura T, Shibata S, Ichikawa Y, Kimura T, et al. Mutations of pre-core and basal core promoter before and after hepatitis B e antigen seroconversion. World J Gastroenterol. 2015;21(2):541–8. Epub 2015/01/17. doi:10.3748/wjg.v21.i2.541. PubMed PMID:25593470; PubMed Central PMCID:PMCPMC4292286.

22. Alexopoulou A, Karayiannis P. HBeAg negative variants and their role in the natural history of chronic hepatitis B virus infection. World J Gastroenterol. 2014;20(24):7644–52. Epub 2014/07/01. doi:10.3748/wjg.v20.i24.7644. PubMed PMID:24976702; PubMed Central PMCID:PMCPMC4069293.

23. Wang XL, Ren JP, Wang XQ, Wang XH, Yang SF, Xiong Y. Mutations in pre-core and basic core promoter regions of hepatitis B virus in chronic hepatitis B patients. World J Gastroenterol. 2016;22(11):3268–74. Epub 2016/03/24. doi:10.3748/wjg.v22.i11.3268. PubMed PMID:27004005; PubMed Central PMCID:PMCPMC4790003.

24. Liu Y, Zhao ZH, Lv XQ, Tang YW, Cao M, Xiang Q, et al. Precise analysis of the effect of basal core promoter/precore mutations on the main phenotype of chronic hepatitis B in mouse models. J Med Virol. 2020. Epub 2020/05/20. doi:10.1002/jmv.26025. PubMed PMID:32427358.

25. Cao M, Zhao Z, Tang Y, Wei Q, Wang L, Xiang Q, et al. A new hepatitis B virus e antigen-negative strain gene used as a reference sequence in an animal model. Biochem Biophys Res Commun. 2018;496(2):502–7. Epub 2018/01/18. doi:10.1016/j.bbrc.2018.01.081. PubMed PMID:29339154.

26. Milich DR, Schodel F, Hughes JL, Jones JE, Peterson DL. The hepatitis B virus core and e antigens elicit different Th cell subsets: antigen structure can affect Th cell phenotype. J Virol. 1997;71(3):2192–201. Epub 1997/03/01. doi:10.1128/JVI.71.3.2192-2201.1997. PubMed PMID:9032353; PubMed Central PMCID:PMCPMC191326.

27. Milich DR, Chen MK, Hughes JL, Jones JE. The secreted hepatitis B precore antigen can modulate the immune response to the nucleocapsid: a mechanism for persistence. J Immunol. 1998;160(4):2013–21. Epub 1998/02/20. PubMed PMID:9469465.

28. Tian Y, Kuo CF, Akbari O, Ou JH. Maternal-Derived Hepatitis B Virus e Antigen Alters Macrophage Function in Offspring to Drive Viral Persistence after Vertical Transmission. Immunity. 2016;44(5):1204–14. Epub 2016/05/10. doi:10.1016/j.immuni.2016.04.008. PubMed PMID:27156385; PubMed Central PMCID:PMCPMC4871724.

29. Xu D, Fu J, Jin L, Zhang H, Zhou C, Zou Z, et al. Circulating and liver resident CD4+CD25+ regulatory T cells actively influence the antiviral immune response and disease progression in patients with hepatitis B. J Immunol. 2006;177(1):739–47. Epub 2006/06/21. doi:10.4049/jimmunol.177.1.739. PubMed PMID:16785573.

30. Dietze KK, Schimmer S, Kretzmer F, Wang J, Lin Y, Huang X, et al. Characterization of the Treg Response in the Hepatitis B Virus Hydrodynamic Injection Mouse Model. PLoS One. 2016;11(3):e0151717. Epub 2016/03/18. doi:10.1371/journal.pone.0151717. PubMed PMID:26986976; PubMed Central PMCID:PMCPMC4795771.

